# The GPR4 antagonist NE-52-QQ57 increases survival, mitigates the hyperinflammatory response and reduces viral load in SARS-CoV-2-infected K18-hACE2 transgenic mice

**DOI:** 10.1101/2024.12.26.630404

**Authors:** Xin-Jun Wu, Karen A. Oppelt, Ming Fan, Mona A. Marie, Madison M. Swyers, Ashley J. Williams, Isabelle M. Lemasson, Rachel L. Roper, Paul Bolin, Li V. Yang

**Author notes:** To whom correspondence should be addressed, Li V. Yang, PhD, Department of Internal Medicine, Brody School of Medicine, East Carolina University 600 Moye Blvd. Greenville, NC 27834, USA.

## Abstract

COVID-19 (Coronavirus disease 19) is caused by infection with SARS-CoV-2 (severe acute respiratory syndrome coronavirus 2) in the respiratory system and other organ systems. Tissue injuries resulting from viral infection and host hyperinflammatory responses may lead to moderate to severe pneumonia, systemic complications, and even death. While anti-inflammatory agents have been used to treat patients with severe COVID-19, their therapeutic effects are limited. GPR4 (G protein-coupled receptor 4) is a pro-inflammatory receptor expressed on vascular endothelial cells, regulating leukocyte infiltration and inflammatory responses. In this study, we evaluated the effects of a GPR4 antagonist, NE-52-QQ57, in the SARS-CoV-2-infected K18-hACE2 transgenic mouse model. Our results demonstrated that GPR4 antagonist treatment increased the survival rate in this severe COVID-19 mouse model. The inflammatory response, characterized by proinflammatory cytokines and chemokines, was reduced in the GPR4 antagonist group compared with the vehicle group. Additionally, both SARS-CoV-2 RNA copy numbers and infectious viral titers in the mouse lung were decreased in the GPR4 antagonist group. The percentage of SARS-CoV-2 antigen-positive mouse brains was also decreased in the GPR4 antagonist group compared to the vehicle group. Furthermore, the GPR4 antagonist inhibited SARS-CoV-2 propagation in Vero E6 cells. Together, these results suggest that GPR4 antagonism may be explored as a novel approach for the treatment of COVID-19 and other similar viral diseases.

## Introduction

Infection with SARS-CoV-2 (severe acute respiratory syndrome coronavirus 2) can cause injuries to the lung and other organs, resulting in the development of COVID-19 (Coronavirus disease 19) disease symptoms in patients. The clinical features of COVID-19 vary widely, ranging from asymptomatic infection, mild illness, to severe disease with respiratory failure and death (1, 2). Inflammatory cell infiltration, diffuse alveolar damage, edema, and thromboembolism in the lung are major pulmonary pathological findings associated with severe COVID-19. Additionally, extrapulmonary manifestations and multiple organ dysfunctions have been observed in COVID- 19 patients (3–5). Autopsy studies on patients succumbing to COVID-19 demonstrate that in addition to the respiratory system, SARS-CoV-2 is widely distributed in multiple anatomic sites including the brain, heart, lymph node, gastrointestinal tract, and other tissues (6).

To date, several therapeutic modalities have been developed, including antivirals, anti-inflammatory agents, and immunomodulators, to treat COVID-19 patients (7, 8). As shown in a randomized clinical trial, treatment with the antiviral drug, remdesivir, accelerated recovery of hospitalized COVID-19 patients and demonstrated a trend of survival benefit (9). Another randomized clinical trial, however, did not observe significant therapeutic benefits of remdesivir in patients with severe COVID-19 (10). Administering PAXLOVID ^TM^ (a combination of nirmatrelvir and ritonavir) to symptomatic COVID-19 patients within 5 days of symptoms reduced the risk of hospitalization or death by 89% compared to a placebo (11). Another study showed that PAXLOVID ^TM^ treatment, versus placebo, did not further alleviate symptoms in patients at standard risk for severe COVID-19 (12).

In addition to antiviral drugs, anti-inflammatory agents have been evaluated for the treatment of COVID-19, as hyperinflammatory responses (“cytokine storm”) are observed in some COVID-19 patients and contribute to poor outcomes (13, 14). SARS-CoV-2 triggered elevated levels of pro-inflammatory cytokines, including interleukin-6 (IL-6) and tumor necrosis factor-alpha (TNF-α) (15), which are associated with pulmonary inflammation and lung damage (16). As shown by the RECOVERY trial, the anti-inflammatory drug, dexamethasone, decreased the mortality rate of hospitalized COVID-19 patients within 28 days (22.9% in the dexamethasone group vs. 25.7% in the usual care group, p < 0.001) (17). Treatment with tocilizumab, a monoclonal antibody targeting the IL-6 receptor, reduced hyperinflammatory responses and provided therapeutic benefits in a subset of COVID-19 patients, but some patients were refractory to tocilizumab treatment (16, 18, 19). Baricitinib is an orally administered inhibitor of Janus kinase 1 (JAK1) and JAK2, exhibiting anti-inflammatory properties. Hospitalized patients were administered a daily dosage of baricitinib, which reduced deaths of COVID-19 by approximately 20% as compared to standard care or placebo (20–23). It remains a significant challenge to effectively treat COVID-19 patients with severe disease to reduce mortality and morbidity.

GPR4 is a pro-inflammatory receptor predominantly expressed in vascular endothelial cells and regulates leukocyte infiltration and inflammatory responses (24–34). Biochemically, GPR4 is partially active at physiological pH and fully activated by extracellular acidic pH (acidosis), which commonly exists in inflamed and hypoxic tissues (35–37). Notably, acidosis is a common complication observed in COVID-19 patients with severe disease (38). Activation of GPR4 increases the expression of inflammatory cytokines, chemokines, and adhesion molecules on endothelial cells to facilitate the adhesion and extravasation of leukocytes (24, 25, 27, 29). GPR4 also regulates vascular permeability and exudate formation under inflammatory conditions (33). GPR4 is expressed in various tissues, with high expression in the lung, heart, and kidney (39, 40). The gene expression of GPR4 is up-regulated in COVID-19 patient lung and colon tissues (41, 42). Therefore, we hypothesized that GPR4 plays an integral role in COVID-19 pathophysiology (42). GPR4 antagonists have been shown to reduce inflammation, vessel permeability, exudate formation, angiogenesis, and pain in several preclinical models (25, 26, 28–31, 33).

In this study, we evaluated the effects of a GPR4 antagonist, NE-52-QQ57, in the SARS-CoV-2- infected K18-hACE2 transgenic mouse model. Our results demonstrated that the GPR4 antagonist reduced the inflammatory response and SARS-CoV-2 viral load and increased the survival rate in this severe COVID-19 murine model.

## Materials and Methods

### Animals and ethics statement

The K18-hACE2 transgenic mice, expressing the human angiotensin-converting enzyme 2 (hACE2) under the control of the epithelial cytokeratin 18 (K18) promoter, were purchased from the Jackson Laboratory (JAX, strain # 034860). The K18-hACE2 hemizygous transgenic mice were bred with wild-type (WT) mice in the animal facility of East Carolina University (ECU). The progenies were genotyped using the protocol provided by the Jackson Laboratory. Male and female hemizygous K18-hACE2 mice (approximately 10 months old) were used in the experiments. All SARS-CoV-2 infectivity studies were performed in the ECU animal biosafety level 3 (ABSL-3) laboratory in accordance with a protocol approved by the ECU Institutional Animal Care and Use Committee.

### The SARS-CoV-2 infected K18-hACE2 mouse model and GPR4 antagonist treatment

SARS-CoV-2 was obtained from the Biodefense and Emerging Infections Research Resources Repository (BEI Resources, NR-52281, SARS-Related Coronavirus 2, Isolate USA-WA1/2020). We propagated the SARS-CoV-2 virus in Vero E6 cells and determined viral titer (plaque-forming units, PFU) by plaque assay (43, 44).

The K18-hACE2 transgenic mice were fed standard chow diets and housed individually in ventilated cages to minimize the risk of SARS-CoV-2 cross-infection. To induce COVID-19 in mice, K18-hACE2 transgenic mice were anesthetized with isoflurane and challenged with 1000 PFU SARS-CoV-2 virus in a final volume of 50 µL by pipetting via the intranasal route (45). At 4 dpi (days post-infection), mice showed some body weight loss with respiratory symptoms and were randomly assigned to orally receive either the GPR4 antagonist (NE-52-QQ57, provided by Novartis, 30 mg/kg, q.d.) or the vehicle control (0.5% methylcellulose, 0.5% Tween 80, and 99% water) (12 mice per group, 6 males and 6 females, from a total of 3 independent experiments). The effects on COVID-19 disease severity and body weight were monitored daily. Animals were weighed daily throughout the study period. The mice were euthanized either upon meeting the criteria for humane endpoints (weight loss exceeding 20% of their initial body weight or displaying severe clinical signs such as lethargy, inactivity, hunched posture, and/or markedly increased or decreased respirations) or at the end of the experiment, which occurred at 10 dpi. Additionally, age and sex-matched K18-hACE2 mice were anesthetized with isoflurane and intranasally administrated 50 µL phosphate-buffered saline (PBS) and euthanized 10 days after the PBS administration as the baseline control.

### Sample processing

Mice were euthanized via intraperitoneal (IP) injection of 250 mg/kg of TBE (tribromoethanol). Blood was collected from the mouse heart and left at room temperature for 20 minutes to form a clot. Then, the tube was centrifuged at 2000g for 10 minutes to separate the serum from the cellular components. The serum was immediately frozen at -80 °C for later cytokine analysis. Mouse right lungs were harvested and weighed. The right lungs were homogenized in 1 ml of FBS (fetal bovine serum)-free Dulbecco’s Modified Eagle Medium (DMEM) using ceramic beads (MagNA Lyser Green Beads, Roche, #03358941001) in a MagNA Lyser instrument (Roche Life Science) for 30s. The homogenates were stored at -80 °C. The left lungs and brains were fixed in 10% neutral-buffered formalin for histology.

### Histology and immunohistochemistry (IHC)

The tissues were fixed in 10% neutral-buffered formalin for at least one week to inactivate SARS-CoV-2 before further processing. They were then embedded in paraffin and sectioned at a thickness of 5 μm before being stained with hematoxylin and eosin (H&E). Lung slides were scored histopathologically based on edema, hemorrhage, alveoli collapse and alveolar wall thickness, graded as 0, 1, 2, 3 which corresponded to healthy, slight, moderate, and severe levels of lung histopathology, respectively.

Additionally, lung and brain slides were immunostained with anti-CD4 (Abcam, #ab183685, dilution 1:500), anti-CD8 (Abcam, #ab217344, dilution 1:500), or SARS-CoV-2 specific anti-nucleocapsid antibody (Novus Biologicals, #NB100-56683, dilution 1:500) and horseradish peroxidase (HRP) conjugated anti-rabbit secondary antibody following the manufacturer’s instructions. Briefly, sections were deparaffinized, and antigen retrieval was performed. Endogenous peroxidase blocking was carried out by treating slides with H_2_O_2_. Endogenous biotin, biotin receptors, and avidin binding sites were blocked with an Avidin-biotin blocking kit (Life Technologies, #004303), followed by a subsequent normal serum blocking step. Slides were incubated with the primary antibody overnight at 4 °C and processed using the VECTASTAIN® Elite ABC-HRP Rabbit IgG Kit (Vector Laboratories, #PK-6101) according to the manufacturer’s instructions. HRP detection was achieved through incubation with the secondary antibody followed by DAB (3,30-diaminobenzidine, Vector Laboratories, #SK-4105) development for 5 minutes. Tissues were counterstained with Gill’s hematoxylin. As a negative control, lung and brain slides from three uninfected mice treated with PBS were evaluated for comparison. The slides were then scanned using an optical microscope (Zeiss Axio Imager M2 microscope) with a 10× objective to check the existence of positive staining of SARS-CoV-2 antigens.

### Plaque assay

To compare viral titers of the lung homogenates, plaque assays in Vero E6 cells were performed. Lung lysate samples were serially diluted ten-fold with serum-free DMEM medium and used to infect Vero E6 cells seeded in a 12-well plate (1.5 × 10^5^ cells per well). The plate was then incubated at 37 °C in a 5% CO_2_ atmosphere for 1 hour with intermittent rocking every 10 minutes to prevent cellular desiccation and facilitate viral binding. Following this viral adsorption, the medium was removed and cells were overlaid with 1 mL of a mixture containing DMEM, 2% fetal bovine serum (FBS), 1% Penicillin/Streptomycin, and 0.4% methylcellulose. After three days of incubation, the methylcellulose overlays were gently removed, and the cells were fixed with 10% neutral-buffered formalin for 20 min. Subsequently, the 10% neutral-buffered formalin solution was removed, and 0.05% (w/v) crystal violet stain solution in distilled water was added for 10 minutes. Viral plaques were counted, and viral titers were calculated as plaques/mg lung.

### Quantitative reverse transcription-polymerase chain reaction (RT-qPCR)

Total RNA was isolated using the DNA/RNA/Protein extraction kit (IBI Scientific, #IB47702), and cDNA was synthesized using 1 μg of RNA (SuperScript IV First Strand cDNA synthesis Reaction, Invitrogen, #18091050). Subsequently, RT-qPCR was performed using a Quantstudio 3 Flex real-time PCR machine with 2× TaqMan Universal Master Mix II (Applied Biosystems, #2704844). Commercial primers/probe sets specific for mouse *Gpr4* (Mm01322176_s1), *Il-1β* (Mm00434228_m1), *Il-6* (Mm00446190_m1), *Il-10* (Mm01288386_m1), *Il-18* (Mm00434226_m1), *Tnf-α* (Mm00443258_m1), *Atf3* (Mm00476033_m1), *Cox2/Ptgs2* (Mm00478374_m1), *Cxcl2* (Mm00436450_m1), *E-selectin* (Mm00441278_m1), *Icam1* (Mm00516023_m1), *Vcam1* (Mm01320970_m1), mouse *Ace2* (Mm01159006_m1), human *ACE2* (Hs01085333_m1), mouse *Tmprss2* (Mm00443687_m1), and human *TMPRSS2* (Hs01122322_m1) were purchased from ThermoFisher and the expressions were normalized to *18S* rRNA (HS99999901_s1) levels.

Additionally, RT-qPCR of the SARS-CoV-2 RNA copy number was performed to test the viral load of lung tissue using the primer kit from Biosearch Technologies (2019-nCoV CDC probe and primer kit for SARS-CoV-2, KIT-nCoV-PP1-1000) targeting the nucleocapsid gene (2019- nCoV_N1): forward primer 5’ GACCCCAAAATCAGCGAAAT 3’, reverse primer 5’ TCTGGTTACTGCCAGTTGAATCTG 3’, and probe 5’ FAM-ACCCCGCATTACGTTTGGTGGACC-BHQ-1 3’. An inactivated SARS-CoV-2 genomic RNA from BEI Resources (#NR-52347) was serially diluted and used as a standard curve control. SARS-CoV-2 levels were determined utilizing a standard curve and expressed as quantified viral RNA copy numbers/µg of RNA.

### Lung and serum cytokines

The lung lysates and serum samples were analyzed using a Luminex 200^TM^ system with a ProcartaPlex Mouse Cytokine/Chemokine Panel 1 26plex platform (ThermoFisher, #EPX260- 26088-901), following the manufacturer’s instructions. Cytokine and chemokine concentrations were determined based on standard curves. Acquired data were analyzed using ProcartaPlex Analysis APP (ThermoFisher). Changes in cytokine levels were analyzed using GraphPad Prism 9 (N=12 for the vehicle group; N=10 for the GPR4 antagonist group as two mice died at night between health checks and their lung and serum samples were not suitable or available for collection for Luminex or RNA analyses).

### *In vitro* assay of the GPR4 antagonist incubated with SARS-CoV-2

GPR4 antagonist (20 μM, 10 μM, 1 μM, and 0.1 μM) or DMSO (dimethyl sulfoxide) vehicle control in 100 μL of DMEM medium were incubated with SARS-CoV-2 (100 PFU, 100 μL) for 1 hour at room temperature, and then the mixture was used to infect Vero E6 cells for 1 hour at 37°C in a 5% CO_2_ atmosphere. Following this incubation, the medium was removed and 1 mL of a mixture containing DMEM, 2% FBS, 1% Penicillin/Streptomycin, and 0.4% methylcellulose was added and incubated for 3 days before plaque counting. For each condition, three biological replicates were used with two experimental repeats.

### Anti-SARS-CoV-2 effects of the GPR4 antagonist post-infection in Vero E6 cells

Confluent Vero E6 cells in 12-well plates were infected with 100 PFU (for RT-qPCR) or 30 PFU (for plaque assay) for 1 hour at 37°C in a 5% CO_2_ incubator. To assess SARS-CoV-2 RNA levels, after 1 hour of virus infection, the culture medium containing the virus was removed. Fresh complete DMEM medium with the GPR4 antagonist or the DMSO vehicle control was added to the cells for a 24-hour treatment. After the treatment, RNA was harvested from Vero E6 cells and RT-qPCR assay was performed to measure the SARS-CoV-2 RNA copy numbers. For the plaque assay, after 1 hour of virus infection, the culture medium containing the virus was removed. Cell medium was changed to DMEM, 2% FBS, 1% Penicillin/Streptomycin, 0.4% methylcellulose, and various concentrations of GPR4 antagonist or the DMSO vehicle control. After 3 days of incubation, the medium was removed, cells were fixed with 10% neutral-buffered formalin, and then stained with crystal violet. For each condition, three biological replicates were used with two experimental repeats.

### Cell viability assay

Cell viability was determined by 3-(4,5-dimethylthiazol-2-yl)-2,5-diphenyltetrazolium bromide (MTT) assay (Roche, #11465007001) and CellTiter-Glo Luminescent Cell Viability (CTG) Assay (Promega, #G7570). Vero E6 cells cultured in 96-well plates (5 × 10^4^ per well) were incubated with the GPR4 antagonist or the DMSO vehicle control for 24 h. For the MTT assay, 20 μL of MTT solution was added per well and incubated at 37 °C for 4 h. After this, the cells were lysed using 200 μL of the solubilization solution and absorbance was measured at 570 nm. For CTG assay, 100 μL of CellTiter-Glo reagent (Promega, #G7573) was added into each well and luminescence intensities were measured using a SpectraMax ID5 plate reader (Molecular Devices). For each condition, three biological replicates were used with two experimental repeats.

### Statistical analysis

Statistical analysis and graph preparation were conducted using GraphPad Prism 9. Results are presented as mean ± SEM (standard error of the mean). The normality of the data was assessed using the Shapiro-Wilk test prior to hypothesis testing. In cases where the data followed a normal distribution, differences between two groups were evaluated using the unpaired t-test. Alternatively, for datasets exhibiting a non-normal distribution, differences between groups were assessed using the Mann-Whitney test. Survival significance was examined using the log-rank (Mantel-Cox) test. The Chi-square test was used to compare the SARS-CoV-2 positive infection rate in the mouse brain. A P value ≤ 0.05 was considered statistically significant.

## Results

### The GPR4 antagonist NE-52-QQ57 increases the survival rate of the K18-hACE2 mice infected with SARS-CoV-2

We intranasally inoculated 1000 PFU of SARS-CoV-2 (strain 2019n-CoV/USA_WA1/2020) into 10-month-old male and female hemizygous K18-hACE2 transgenic mice. K18-hACE2 mice infected with SARS-CoV-2 developed severe disease. These mice exhibited weight loss of 10- 20%, lethargy, inactivity, ruffled fur, markedly increased or decreased respirations, and hunched posture, with most mice reaching the humane endpoint or dying by 8 dpi. Starting from 4 dpi, we administered either vehicle or the GPR4 antagonist NE-52-QQ57 to assess its effects on SARS-CoV-2-induced inflammatory responses and disease severity in the K18-hACE2 mouse model.

Our results demonstrated that 25% of the mice in the vehicle control group (group size equals 6 males and 6 females) survived at 10 dpi. In comparison to vehicle controls, 66.7% of the mice in the GPR4 antagonist-treated group (group size equals 6 males and 6 females) survived at 10 dpi. The results indicate that the GPR4 antagonist treatment increased the survival rate in this severe COVID-19 mouse model (p = 0.03, by log-rank (Mantel-Cox) test, Fig.1A). All infected mice were weighed daily throughout the study period. Beginning at 5 dpi, all SARS-CoV-2-infected K18-hACE2 mice demonstrated marked weight loss. GPR4 antagonist treatment decreased the amount of weight loss, but differences between vehicle and treatment groups were not statistically significant (Fig. 1B). Notably, all surviving mice started regaining weight after 7-8 dpi (Fig. 1B) and their overall health condition and activity were gradually improved by 10 dpi.

**Figure 1.**
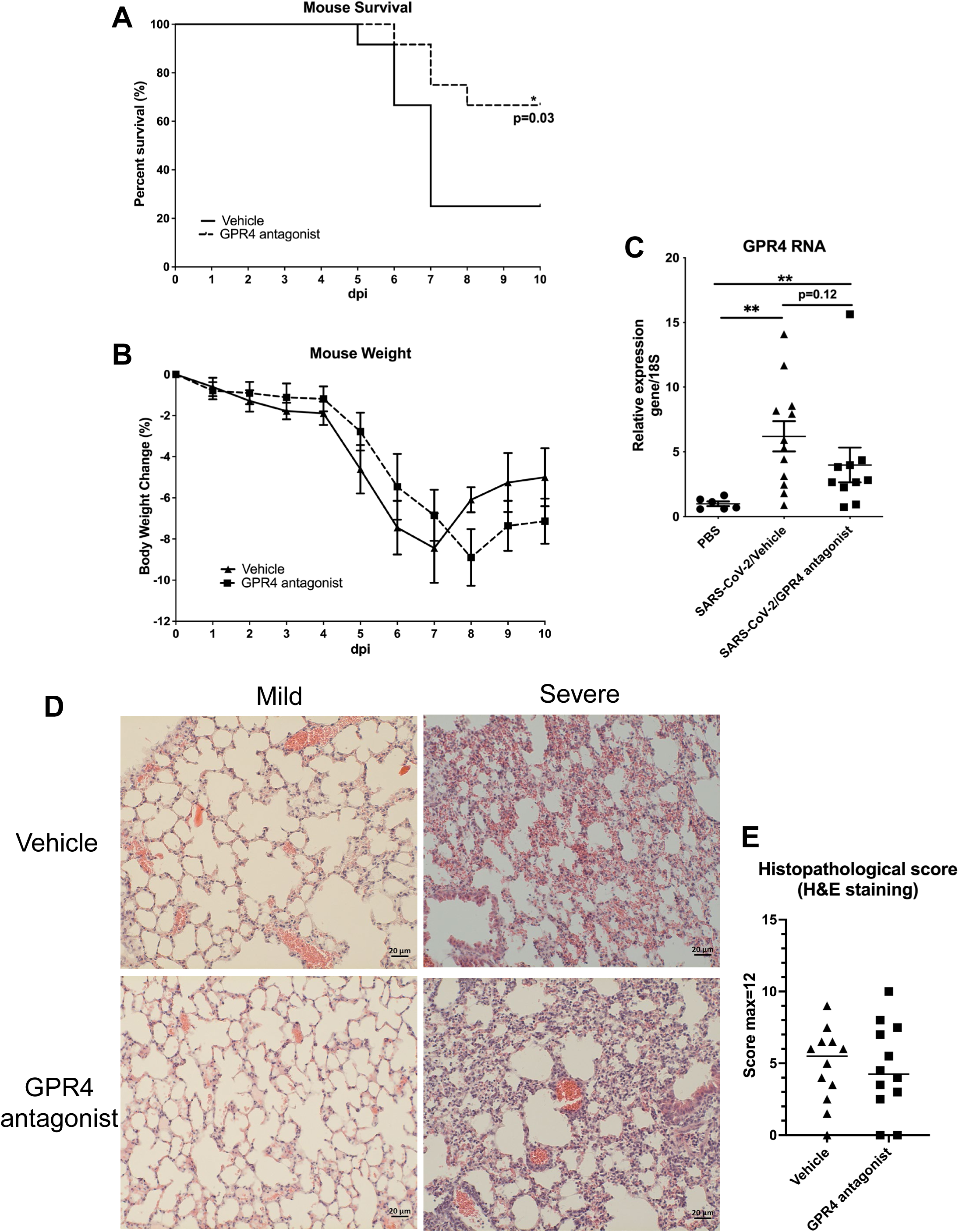
Treatment with GPR4 antagonist NE-52-QQ57 improves survival of SARS-CoV-2-infected K18-hACE2 mice. Mice were treated with GPR4 antagonist NE-52-QQ57 or vehicle control for up to 6 days starting from 4 dpi. (**A)** The survival rate of SARS-CoV-2-infected K18-hACE2 mice is increased by the administration of the GPR4 antagonist. Ten-month-old male and female K18-hACE2 transgenic mice were intranasally inoculated with 1000 PFU of SARS-CoV-2 (N=12). Survival analysis was performed using the Kaplan-Meier method with a log-rank (Mantel-Cox) test, * p < 0.05. (**B)** Daily body weight changes in GPR4 antagonist-treated or vehicle control mice were recorded up to 10 dpi or until the mice reached the humane endpoint. The difference in body weight change was analyzed using multiple unpaired t-tests. (**C)** RT-qPCR was conducted to quantify the expression of GPR4 in non-infected (PBS) and SARS-CoV-2-infected mouse lung tissues (compared using two-tailed Mann-Whitney test) (N=6 for PBS no virus inoculation; N=12 for vehicle, N=10 for GPR4 antagonist). Error bars indicate means ± SEM. ** p < 0.01. **(D)** Representative pictures of mouse lung histology (H&E staining) with mild or severe histopathology in vehicle or GPR4 antagonist-treated mice. Scale bar = 20 µm. **(E)** Mouse lung histopathological score. Two-tailed Student’s t-test did not indicate significance.

### SARS-CoV-2 induced GPR4 expression in the mouse lungs

Quantitative RT-PCR was performed to examine GPR4 RNA expression in the mouse lungs infected with SARS-CoV-2 and the control mouse lungs. GPR4 RNA expression levels were significantly increased by about 7-fold in the lungs of K18-hACE2 mice infected with SARS-CoV-2 when compared to the lungs of control mice intranasally administrated with PBS (Fig. 1C). Within the mice infected with SARS-CoV-2, a trend of reduced GPR4 expression was observed in the GPR4 antagonist group compared to the vehicle group (Fig. 1C). For lung histological analysis, there was a trend (not statistically significant) of reduced histopathological scores of SARS-CoV-2 infected mouse lung tissues in the GPR4 antagonist group compared with the vehicle control group (Fig. 1D and 1E). Additionally, 1 out of 12 mice in the vehicle control SARS-CoV- 2 infected group had brain hemorrhage which was not observed in the GPR4 antagonist group (Supplementary Fig. 1).

### Reduced pro-inflammatory cytokine and chemokine levels in the GPR4 antagonist-treated K18-hACE2 mice infected with SARS-CoV-2

COVID-19 disease progression is correlated with significant alterations in cytokine profiles (16, 46). To evaluate the impact of GPR4 antagonist treatment on the inflammatory response to SARS-CoV-2 infection, RT-qPCR was employed to analyze the RNA levels of inflammatory genes in the lung tissues of SARS-CoV-2-infected K18-hACE2 mice. Compared with the lungs of vehicle control mice, GPR4 antagonist treatment significantly reduced the expression of the inflammatory genes *Cxcl2*, *Ptgs2*, and *Icam1* in the lungs of K18-hACE2 mice infected with SARS-CoV-2 (Fig. 2A). Moreover, other inflammatory genes (e.g. *Il-1β*, *Il-6*, *Il-10*, *Il-18*, *Tnf-α*, *E-selectin*, *Atf3*, and *Vcam1*) also showed a strong trend of reduced expression, although not statistically significant (Fig. 2A). In addition, cytokine and chemokine protein levels in the lung tissue and serum were measured using a mouse 26 plex Luminex panel (Fig. 2B, 2C, Supplementary Fig. 2 and 3). Similar to the RT-qPCR result, CXCL2 protein showed a trend of reduced expression in the mouse lung in the GPR4 antagonist group (Fig. 2B). Furthermore, there was a seven-fold decrease of IL6 levels in the GPR4 antagonist group (26.48±3.956 pg/mL) compared with the vehicle group (191±165.8 pg/mL), along with a decrease in IL-10, IL-17A, and CXCL2 levels (Fig. 2B). Regarding serum cytokines and chemokines, the T cell-associated cytokine IFNγ showed a significant decrease in the GPR4 antagonist group (Fig. 2C). Several other serum cytokines (e.g. IL-1β, IL-6, IL-10, IL- 17A, and TNFα) had a trend of lower expression in the GPR4 antagonist treated mice compared with the vehicle group (Fig. 2C). IL-4 and IL-13 were under detection limits in the serum samples (Supplementary Fig. 3). Overall, our data suggest that the GPR4 antagonist treatment dampens the expression levels of cytokines and chemokines induced by SARS-CoV-2 infection.

**Figure 2.**
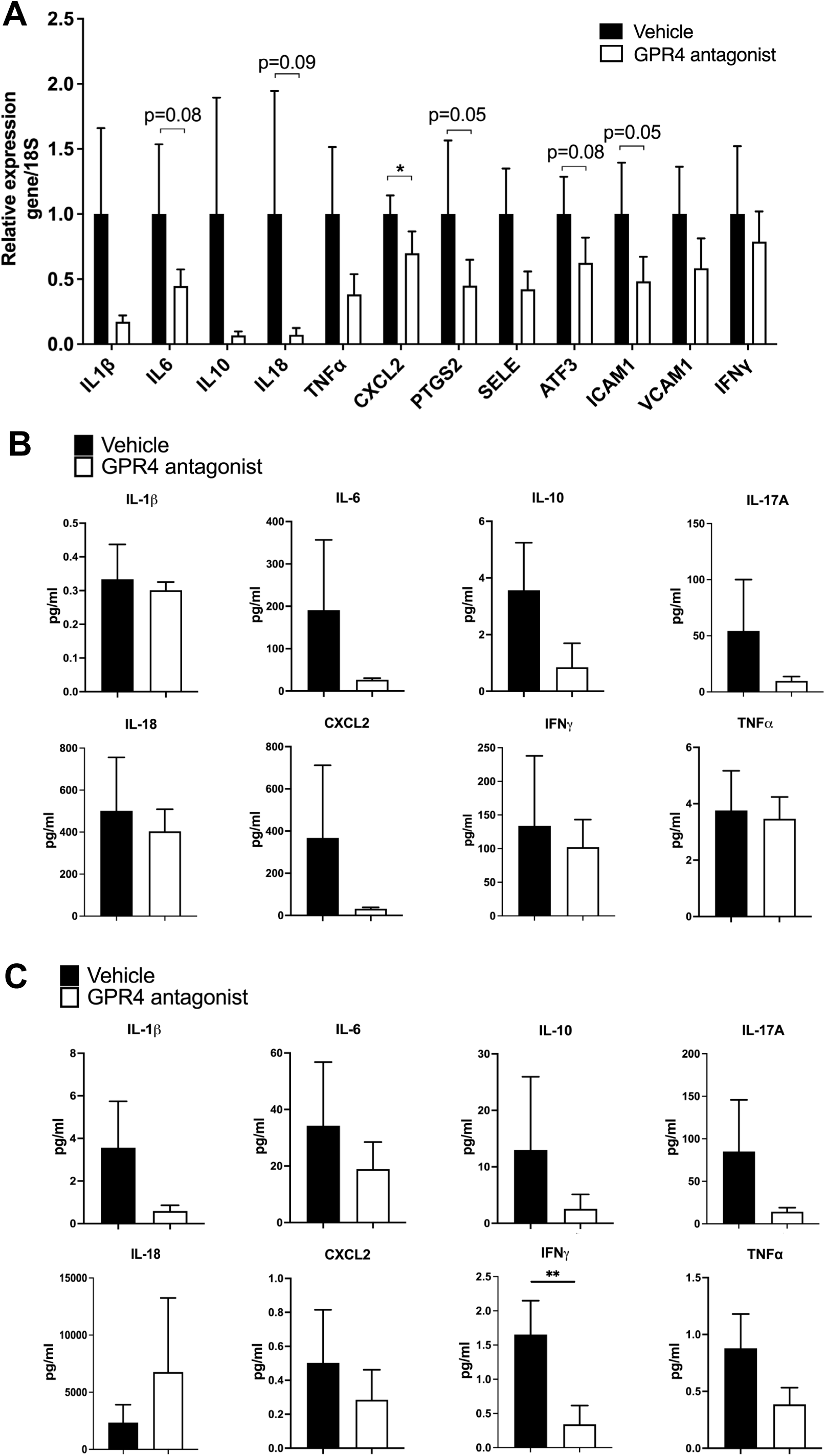
GPR4 antagonist treatment reduces cytokine and chemokine levels in K18-hACE2 mice infected with SARS-CoV-2. **(A)** Fold change in gene expression levels of specified cytokines, chemokines, and other inflammatory genes assessed via RT-qPCR and normalized to 18S rRNA, compared with vehicle controls in mouse lung homogenates (N=12 for vehicle, N=10 for GPR4 antagonist). * p < 0.05. **(B)** Cytokine/chemokine protein levels in mouse lung tissues measured by the Luminex multiplex platform. **(C)** Cytokine/chemokine protein levels in mouse serum measured by the Luminex multiplex platform. Statistical differences in cytokine/chemokine levels were analyzed using the one-tailed Mann-Whitney test (N=12 for vehicle, N=10 for GPR4 antagonist). Error bars indicate mean ± SEM.

### GPR4 antagonist treatment decreases the viral loads in the lungs and brains of the K18- hACE2 mice infected with SARS-CoV-2

To assess the viral load, mouse lung lysates were analyzed by RT-qPCR and plaque assays. Treatment with the GPR4 antagonist reduced SARS-CoV-2 viral loads in the mouse lung tissues. RT-qPCR revealed that SARS-CoV-2 viral RNA copy numbers were reduced about 11.4-fold in the lungs of GPR4 antagonist-treated mice compared to the vehicle treatment group (6.28 × 10^8^ ± 3.99 × 10^8^ copies/μg RNA vs. 7.18 × 10^9^ ± 3.20 × 10^9^ copies/μg RNA, p = 0.08; Fig. 3A). Likewise, plaque assays demonstrated a statistically significant reduction in infectious SARS-CoV-2 in the mouse lung tissues of the GPR4 antagonist group compared to the vehicle-control group (73.83 ± 63.50 pfu/mg vs. 397.6 ± 213.0 pfu/mg, p = 0.02; Fig. 3B).

**Figure 3.**
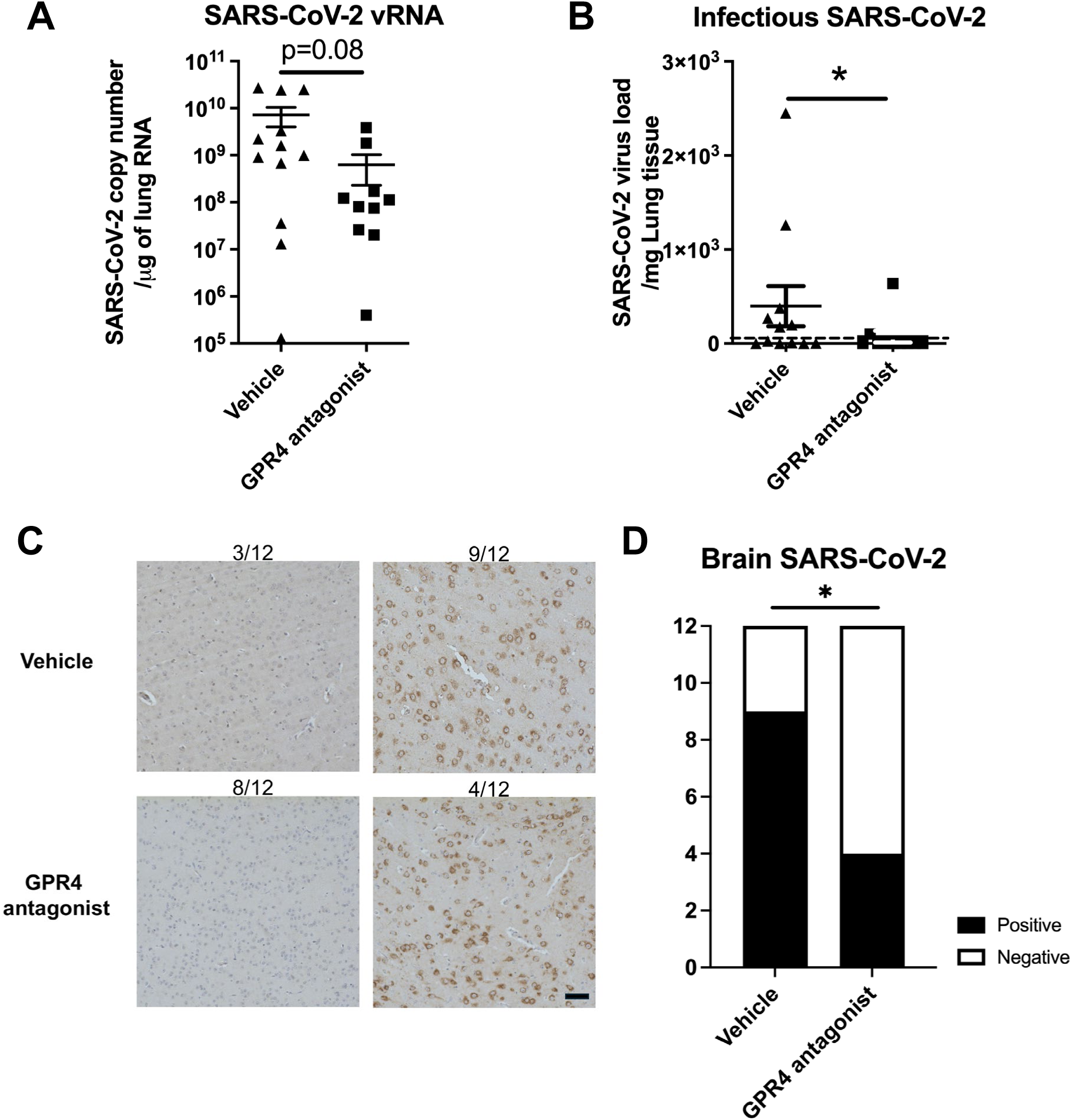
SARS-CoV-2 viral load in the lungs and brains from K18-hACE2 mice that received either GPR4 antagonist or vehicle. **(A)** RT-qPCR to quantify viral RNA levels in mouse lung tissues (RNA copies/μg lung RNA). The data were analyzed using the two-tailed unpaired t-test and shown in mean ± SEM (N=12 for vehicle, N=10 for GPR4 antagonist). **(B)** Plaque assays were analyzed to determine the infectious viral titers (PFU/mg lung) in the lungs of vehicle- and GPR4 antagonist-treated mice infected with SARS-CoV-2. The limit of detection (LOD = 5 PFU/mg lung) is indicated by the dotted horizontal line. * p < 0.05. **(C)** Analysis of SARS-CoV-2 virus nucleocapsid distribution in mouse brain through IHC. The percentage of SARS-CoV-2 positive viral staining in the mouse brain was assessed using a microscope (N=12 for vehicle, N=12 for GPR4 antagonist). Scale bar = 20 μm. Error bars indicate mean ± SEM. **(D)** SARS-CoV-2 positive ratio in the brains of mice treated with GPR4 antagonist or vehicle. Analyzed using the Chi-square test, * p < 0.05.

To evaluate the effects of GPR4 antagonist treatment on viral burden in mouse brains, the presence of the SARS-CoV-2 virus in the brains was assessed through IHC staining with an antibody detecting the viral nucleocapsid protein. Subsequently, the percentage of brain tissue positive for the viral antigen was determined. IHC staining demonstrated that 66.67% of GPR4 antagonist-treated mice (8 out of 12) showed no detection of SARS-CoV-2 in the brain. In contrast, only 25% of the vehicle-treated mice (3 out of 12) showed no detection of SARS-CoV-2 in the brain (Fig. 3C). The GPR4 antagonist group of mice exhibited significantly lower levels of the virus in the brain compared to the vehicle control group (Fig. 3D, Chi-Square test, p = 0.04). All the mice without detectable SARS-CoV-2 by IHC in their brains survived to 10 dpi. Together, these findings demonstrate that GPR4 antagonist treatment systemically reduces SARS-CoV-2 viral loads in K18-hACE2 mice.

### Reduction in CD4^+^ and CD8^+^ immune cell clusters in the brains and lungs of SARS-CoV-2- infected mice treated with the GPR4 antagonist

We further assessed the presence of CD4^+^ and CD8^+^ T cells in the brains and lungs of SARS-CoV- 2-infected mice by IHC, as these cells are essential for the inflammatory and immune responses. CD4^+^ and CD8^+^ T cell infiltration was observed on stained sections as clusters or scatterings of cells in the brains of SARS-CoV-2-infected mice (Fig. 4A and 4C). Compared to the vehicle group, GPR4 antagonist-treated mice exhibited fewer CD4^+^ and CD8^+^ immune cell clusters in the mouse brain sections (Fig. 4B and 4D). Moreover, fewer CD4^+^ and CD8^+^ T cell clusters were detected in the lung sections of SARS-CoV-2-infected mice treated with the GPR4 antagonist (Supplementary Fig. 4).

**Figure 4.**
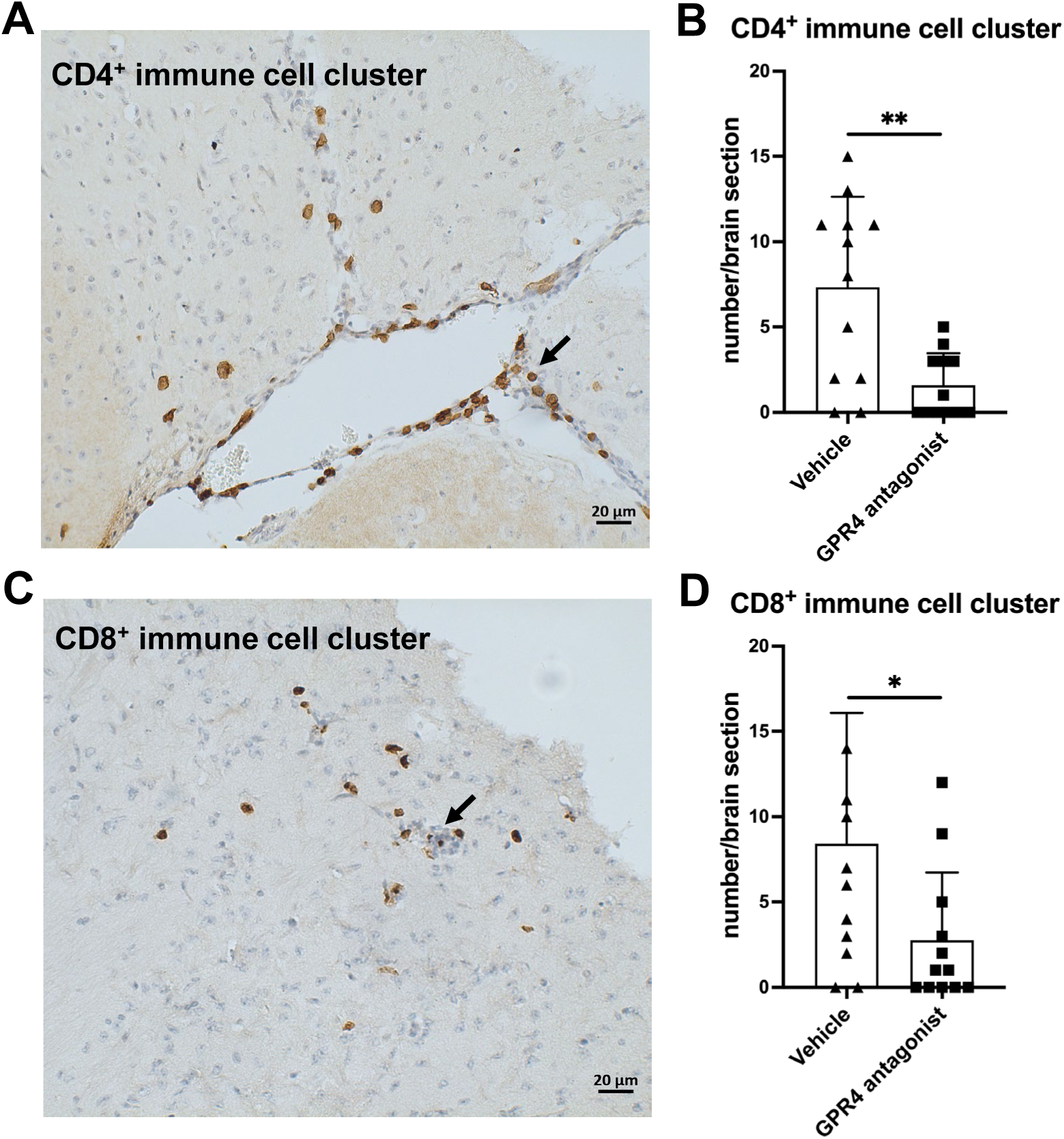
GPR4 antagonist treatment reduces CD4^+^ and CD8^+^ immune cell clusters in the brains of SARS-CoV-2-infected K18-hACE2 mice. **(A)** A representative image of a CD4^+^ immune cell cluster in the mouse brain, visualized using IHC with an antibody detecting CD4. Scale bar = 20 μm. **(B)** Quantification of the number of CD4^+^ immune cell clusters in the brain using microscopy. Analyzed using the two-tailed Student’s t-test, ** p < 0.01. Error bars represent mean ± SEM. N=12 for the vehicle group, and N=12 for the GPR4 antagonist group. **(C)** A representative image of a CD8^+^ immune cell cluster in the mouse brain, visualized using IHC with an antibody detecting CD8. Scale bar = 20 μm. **(D)** Quantification of the number of CD8^+^ immune cell clusters in the brain using microscopy. Analyzed by the two-tailed Student’s t-test, * p < 0.05. Error bars represent mean ± SEM. N=12 for the vehicle group, and N=12 for the GPR4 antagonist group.

### GPR4 antagonist inhibits SARS-CoV-2 propagation *in vitro*

As the GPR4 antagonist treatment reduced the viral load in the lungs and brains of K18-hACE2 mice infected with SARS-CoV-2 (Fig. 3), we then used cell cultures to assess the potential anti- viral effects of the GPR4 antagonist in pharmacologically relevant concentrations as previously demonstrated in mouse models (30). To investigate if the GPR4 antagonist could inactivate SARS-CoV-2 directly, we incubated SARS-CoV-2 (100 PFU) with the GPR4 antagonist (20 μM, 10 μM, 1 μM, and 0.1 μM) or the DMSO vehicle control for 1 hour at room temperature before infecting Vero E6 cells. Plaque assays demonstrated that the GPR4 antagonist could not directly inactivate SARS-CoV-2, as similar plaque numbers were observed in the GPR4 antagonist groups compared with the DMSO group (Fig. 5A). To determine if the GPR4 antagonist had anti-SARS-CoV-2 effects when administered post-infection, we infected Vero E6 cells with virus for a one-hour adsorption period and then treated the cells with the GPR4 antagonist for 24 hours (Fig. 5B). The GPR4 antagonist significantly decreased the RNA copy numbers of SARS-CoV-2 in Vero E6 cells in a dose-dependent manner as assessed by RT-qPCR (Fig. 5B). GPR4 antagonist (20 μM) reduced the genomic RNA levels of SARS-CoV-2 by 8.3-fold (Fig. 5B). Similarly, plaque assays confirmed a decrease in infectious virus after 72 hours of GPR4 antagonist treatment (Fig. 5C). Both CTG assay and MTT assay showed that the GPR4 antagonist (20 μM, 10 μM, 1 μM, 0.1 μM) had no cytotoxic effects on Vero E6 cells (Supplementary Fig. 5).

**Figure 5.**
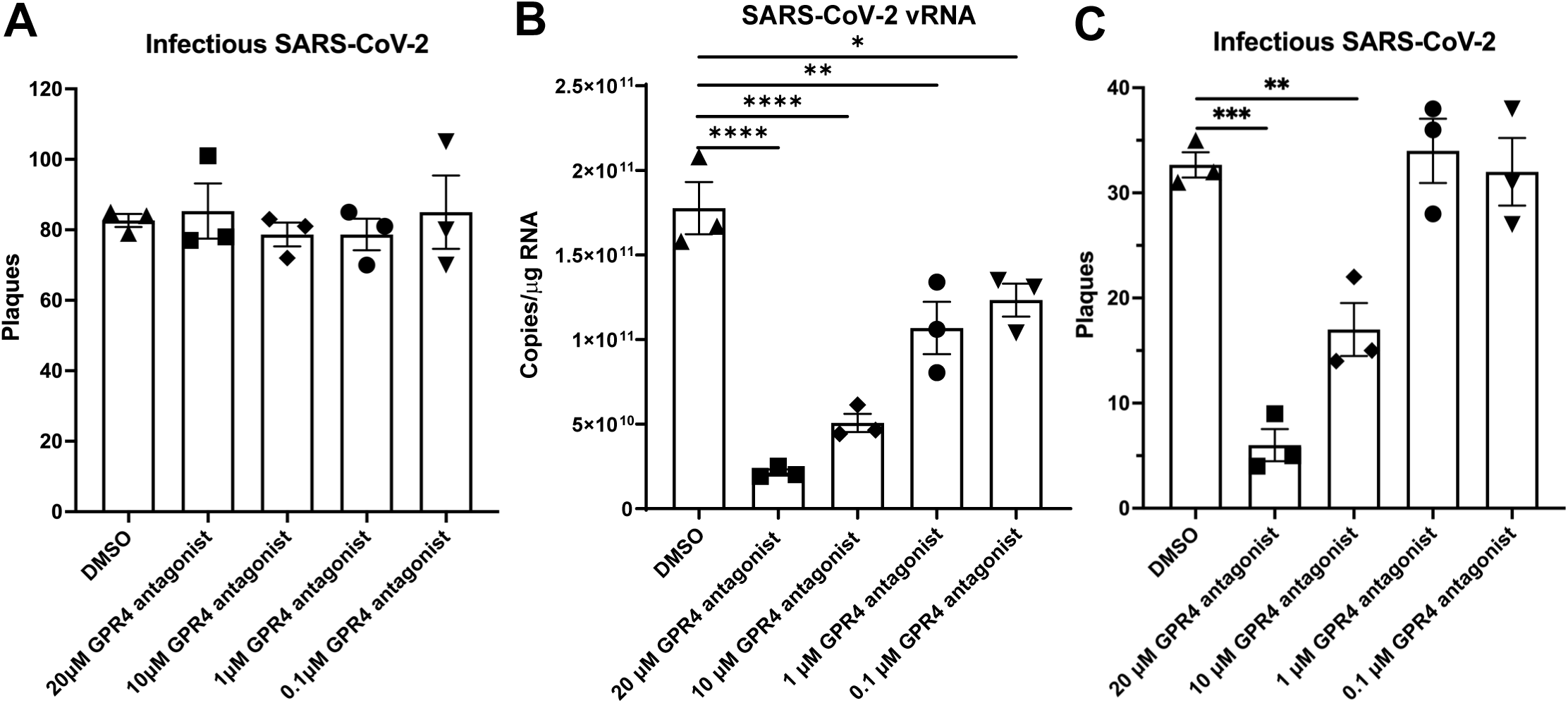
Anti-SARS-CoV-2 effects of GPR4 antagonist *in vitro*. **(A)** GPR4 antagonist incubated with SARS-CoV-2 (100 PFU) for 1h before infecting Vero E6 cells. GPR4 antagonist-containing medium was removed after infection. Plaque formation was measured to determine the infectious SARS-CoV-2 viral titer. N=3 samples. **(B)** Viral RNA in the SARS-CoV-2-infected Vero E6 cells (100 PFU inoculum) treated with various concentrations of GPR4 antagonist was determined 24 h post-infection. The GPR4 antagonist was maintained in the medium until cell assessment 24 h after treatment. Viral RNA isolated from Vero E6 cells was quantified by RT- qPCR targeting the nucleocapsid gene. N=3 samples. * p < 0.05, ** p < 0.01, **** p < 0.0001. **(C)** The infectious viral load by the plaque assay in SARS-CoV-2-infected Vero E6 cells (30 PFU inoculum) in response to treatment of vehicle DMSO or GPR4 antagonist at 72 h post-infection. The GPR4 antagonist was maintained in the medium for 72 h until cell assessment. N=3 samples. ** p < 0.01, *** p < 0.001. Comparisons between groups were analyzed by one-way ANOVA followed by post hoc Dunnett’s test. Error bars indicate mean ± SEM.

## Discussion

Following the onset of the COVID-19 pandemic, vaccines and therapeutic interventions have been developed to address the spectrum of disease severity associated with the SARS-CoV-2 virus. COVID-19 vaccines have been shown to reduce the risk of severe illness, hospitalization, death, and long COVID symptoms (47–49). Antivirals, such as PAXLOVID ^TM^, can also reduce the risk of hospitalization and death in COVID-19 patients (11). Despite the remarkable development of SARS-CoV-2 vaccines and antivirals, some COVID-19 patients may still progress to severe illness, including hospitalization and death. Hyperinflammation and cytokine storm have been demonstrated to play a pivotal role in the pathophysiology of severe COVID-19 and have consistently been linked to an elevated risk of mortality among patients afflicted with the disease (46, 50, 51). Anti-inflammatory and immunosuppressant agents can bring therapeutic benefits to COVID-19 patients with severe disease (7). It has been reported that anti-inflammatory and immunosuppressant agents, such as dexamethasone, tocilizumab, and baricitinib, moderately reduce the mortality rate in hospitalized COVID-19 patients by decreasing the late-phase hyperinflammatory responses(16–23).

In this study, we demonstrated the anti-inflammatory and anti-viral effects of the GPR4 antagonist NE-52-QQ57 in the SARS-CoV-2-infected K18-hACE2 mouse model. Animals administered with the GPR4 antagonist exhibited an increased survival rate compared to those in the vehicle control group in this severe COVID-19 mouse model (Fig. 1A). Also, GPR4 antagonist treatment reduced the expression of inflammatory cytokines, chemokines, adhesion molecules, and PTGS2 (COX2) in the SARS-CoV-2-infected mouse lungs and serum samples (Fig. 2). GPR4 is a pro-inflammatory receptor overexpressed in inflamed tissues (24–34). In line with previous findings that GPR4 expression is upregulated in lung and colon tissues of COVID-19 patients (41, 42), we observed an increase in GPR4 expression in the K18-hACE2 mouse lung following SARS-CoV- 2 infection (Fig. 1C). Altogether, the findings suggest that GPR4 is a pro-inflammatory receptor involved in COVID-19 and GPR4 antagonism may be exploited as a potential therapeutic approach to dampen the hyperinflammatory response of COVID-19 to alleviate disease severity.

In addition to its anti-inflammatory effects, we demonstrated the novel anti-viral effects of the GPR4 antagonist NE-52-QQ57 against SARS-CoV-2. The animals treated with the GPR4 antagonist showed significantly lower viral titers in both lung and brain tissues compared with higher titers in the vehicle control group. These results demonstrate a protective effect of the GPR4 antagonist in the SARS-CoV-2 infection, even though the underlying mechanisms remain unclear. We found that the GPR4 antagonist had no direct effects on inactivating SARS-CoV-2 and did not block SARS-CoV-2 virus entry into Vero E6 cells (Fig. 5A). However, the GPR4 antagonist can inhibit SARS-CoV-2 propagation in Vero E6 cells after the virus enters the cells (Fig. 5B-C). In an attempt to further evaluate potential mechanisms by which the GPR4 antagonist inhibits SARS-CoV-2 propagation, we assessed its effects on the gene expression of ACE2 and TMPRSS2 which are important for viral cellular entry. We found no difference in Vero E6 cells using the TaqMan primer specific for human ACE2 (Supplementary Fig. 6A); however, the TaqMan primer specific for human TMPRSS2 did work with Vero E6 (data not shown). Similarly, no differences in ACE2 or TMPRSS2 expressions were found in A549 human lung epithelial cancer cells, Caco-2 human colon epithelial cancer cells, and mouse Lewis lung carcinoma cells after a 24-hour GPR4 antagonist treatment (Supplementary Fig. 6B-D). These results further support that the GPR4 antagonist does not affect the entry of SARS-CoV-2 into cells (Fig. 5A). As SARS-CoV-2 propagation involves multiple steps such as viral entry, replication, release, and host immune response, further research is needed to elucidate the precise mechanisms by which the GPR4 antagonist reduces SARS-CoV-2 propagation in cells and viral burden in animals.

In addition to the lungs, SARS-CoV-2 can infect the brain and other organs. Neurological complications associated with COVID-19 have the potential to be debilitating and even life-threatening(52–54). Reduced viral burden in mouse brain samples may contribute to the higher survival rate of animals in the GPR4 antagonist-treated group (Fig. 3C-D). Moreover, neurological symptoms are commonly reported in patients with Long Covid (55). SARS-CoV-2 infection in patients is also found to be associated with the rapid progression of pre-existing dementia (56). Future studies are warranted to evaluate the effects of GPR4 antagonism on neurological and other complications related to Long Covid.

In summary, this study highlights the effectiveness of the GPR4 antagonist NE-52-QQ57 in reducing mortality from SARS-CoV-2 infection in K18-hACE2 mice. Animals treated with the GPR4 antagonist exhibited markedly enhanced survival rates, reduced lung and brain SARS-CoV- 2 viral burden, and mitigated inflammatory responses. Our results demonstrate that the GPR4 antagonist NE-52-QQ57 has both anti-inflammatory and anti-viral effects against SARS-CoV-2 infection. Further research is needed to elucidate the mechanisms by which the GPR4 antagonist inhibits SARS-CoV-2 propagation in cells and tissues. It is also crucial to investigate whether the GPR4 antagonist exhibits anti-inflammatory and anti-viral effects in other similar viral diseases in addition to COVID-19.

## Author Contributions

Xin-Jun Wu: Performed the majority of the experiments; Analyzed and interpreted the data; Contributed reagents, materials, analysis tools or data; Wrote the manuscript.

Karen A. Oppelt: Designed and performed the animal experiments; Acquired funding; Edited the manuscript.

Ming Fan, Madison M. Swyers, and Ashley J. Williams: Performed immunohistochemistry. Mona A. Marie: Performed the animal experiments.

Isabelle M. Lemasson: Performed the virus production experiment; Contributed to scientific discussion.

Rachel L. Roper: Contributed resources and scientific discussion; Edited the manuscript. Paul Bolin: Acquired funding; Contributed resources and scientific discussion.

Li V. Yang: Conceived and designed the experiments; Performed the experiments; Analyzed and interpreted the data; Contributed reagents, materials, analysis tools or data; Acquired funding; Supervised the project; Wrote the manuscript.

All the authors reviewed the manuscript.

## Acknowledgments

We thank the ABSL-3 facility staff at East Carolina University (ECU) for technical support, Shayan Nik Akhtar for initial assistance in genotyping K18-hACE2 mice, and Cindy Kukoly in the Department of Internal Medicine and Drs. Gabriel Abuna and Ramiro Murata in the School of Dental Medicine at ECU for their assistance with the Luminex instrument. We also thank Drs. Pius Loetscher and Juraj Velcicky at the Novartis Institutes for BioMedical Research for providing the GPR4 antagonist NE-52-QQ57, and the BEI Resources for providing the SARS-Related Coronavirus 2 (SARS-CoV-2, Isolate USA-WA1/2020).

---

**Supplementary Figure 1.**
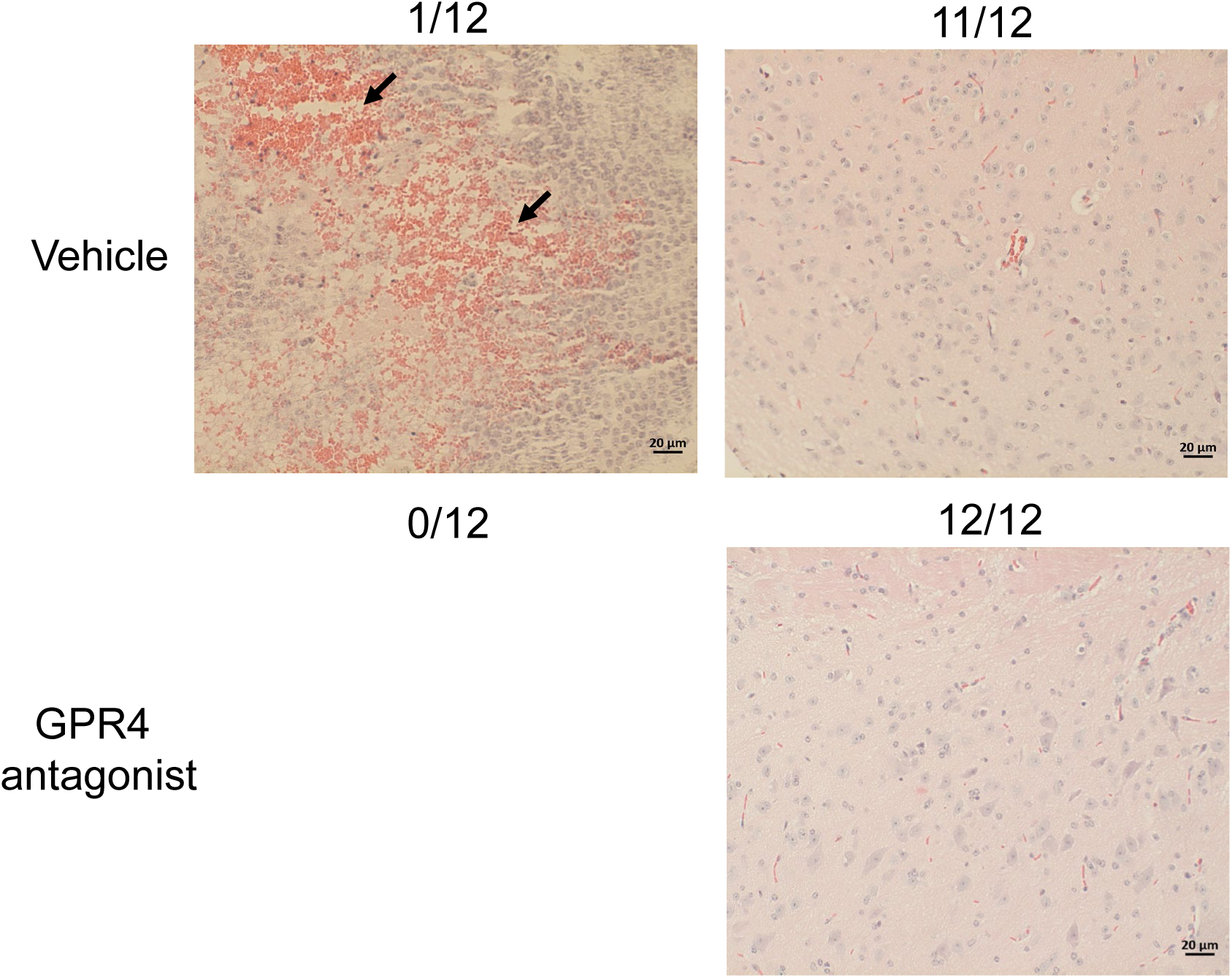
Representative images of H&E staining of mouse brains and their associated hemorrhage rates. Black arrows indicate hemorrhagic areas in the brain of a SARS-CoV-2-infected mouse treated with vehicle. Note no hemorrhagic areas in the brains of mice treated with GPR4 antagonist. Scale bar = 20 μm.

**Supplementary Figure 2.**
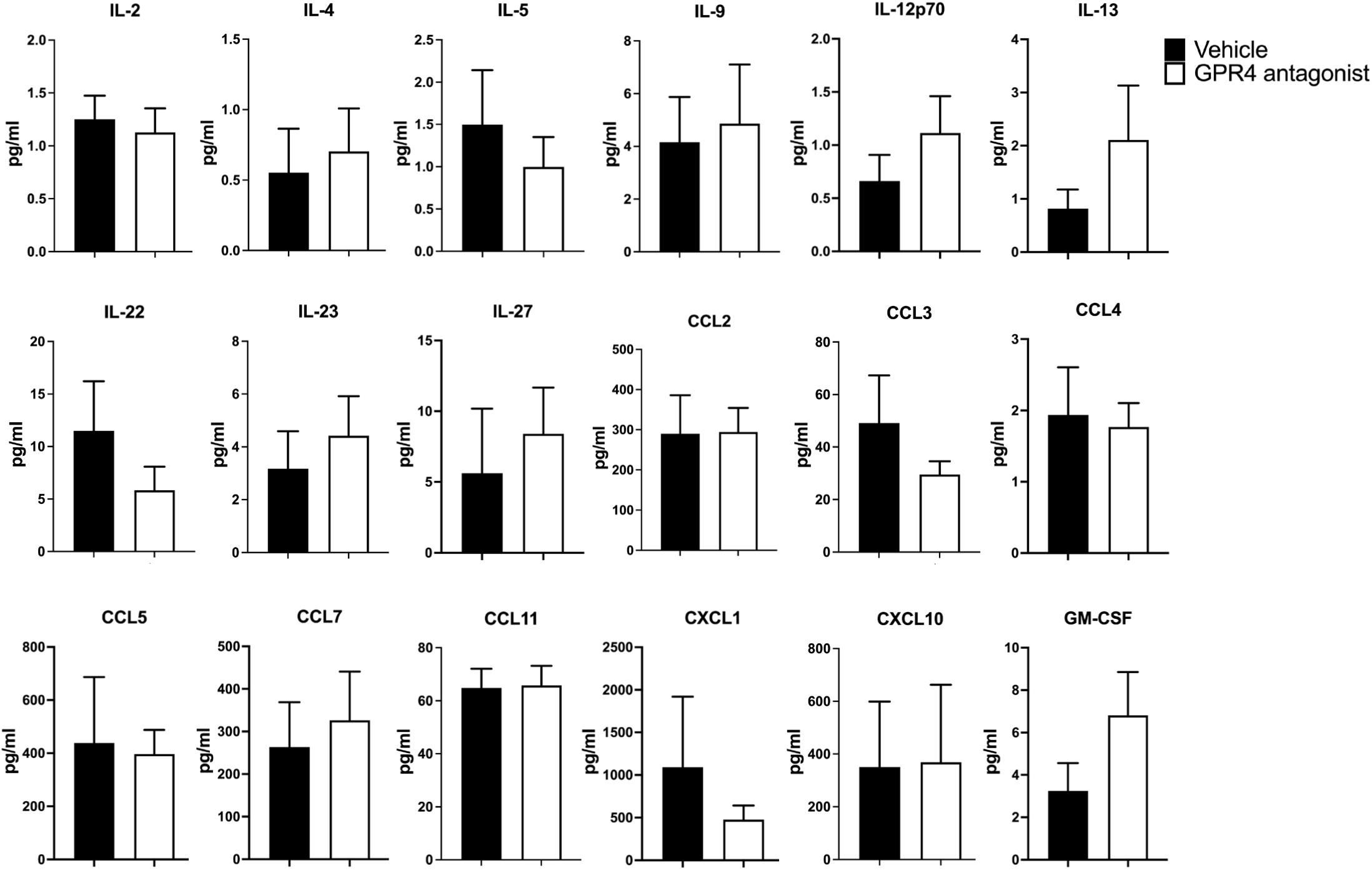
Other cytokines and chemokines in the lung tissues of SARS-CoV-2-infected mice treated with GPR4 antagonist or vehicle. Cytokine/chemokine protein levels in mouse lung tissues were measured by the Luminex multiplex platform.

**Supplementary Figure 3.**
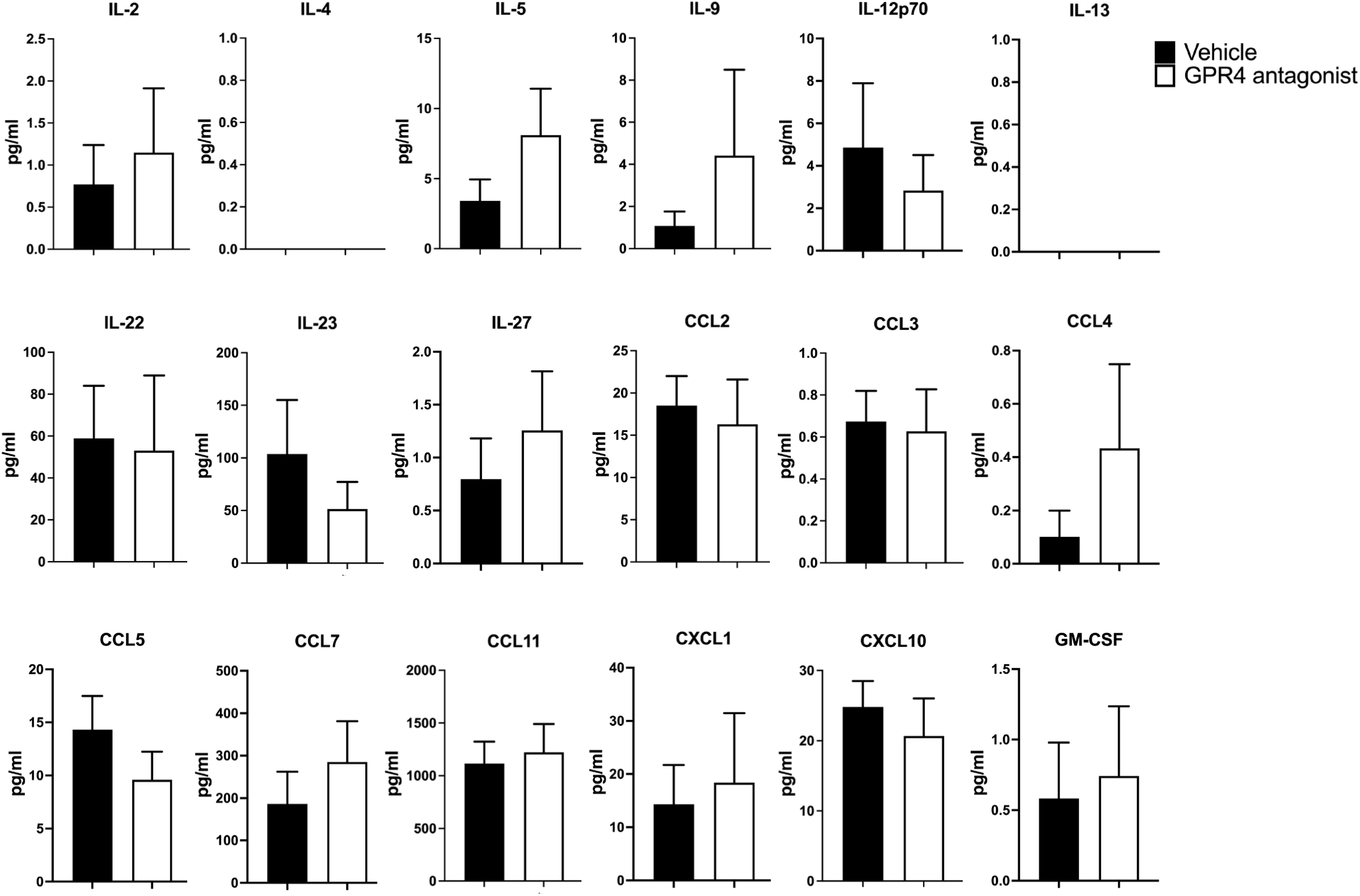
Other cytokines and chemokines in the serum of SARS-CoV-2-infected mice treated with GPR4 antagonist or vehicle. Cytokine/chemokine protein levels in mouse serum were measured by the Luminex multiplex platform.

**Supplementary Figure 4.**
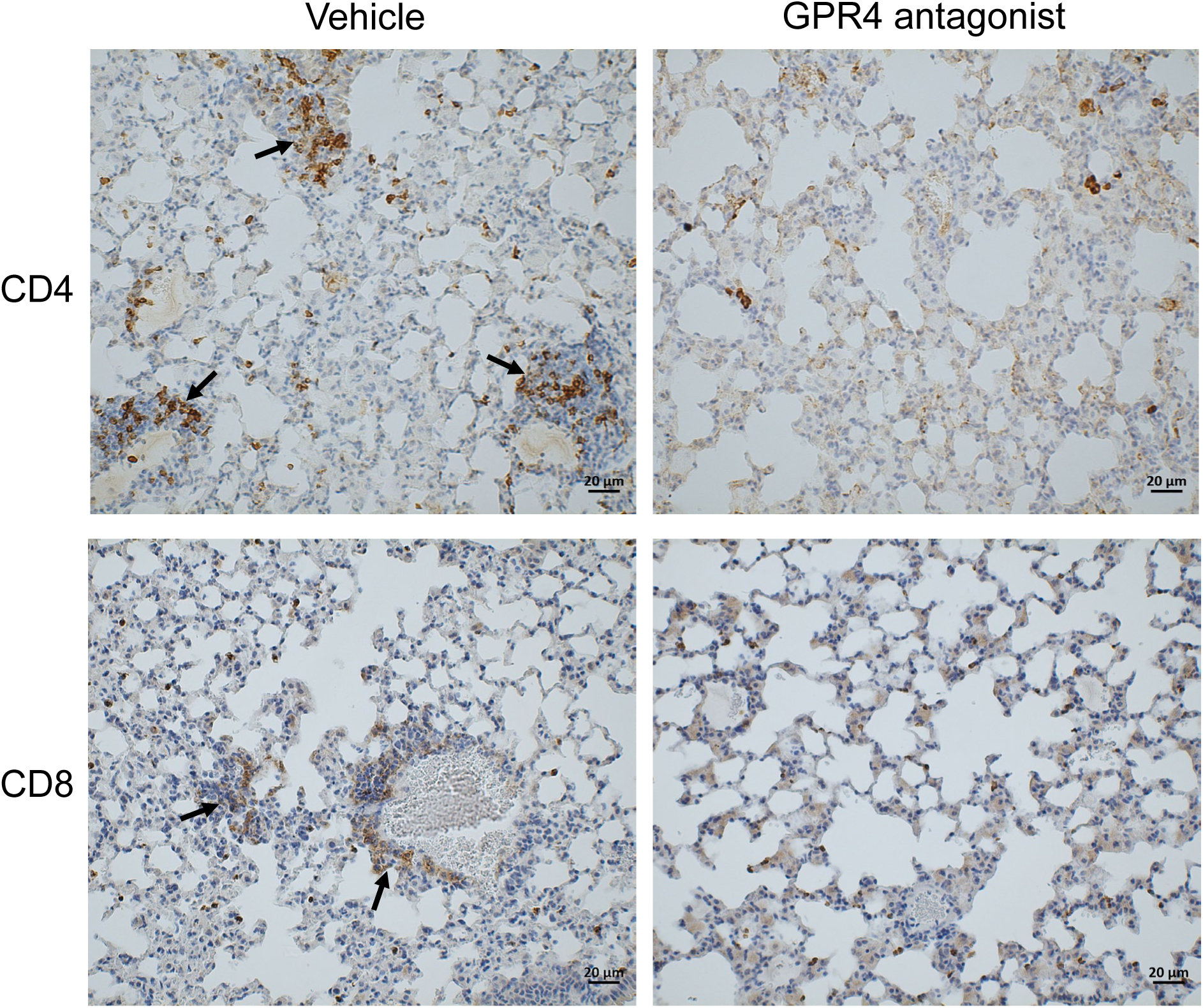
CD4^+^ and CD8^+^ T cell clusters in the mouse lung. GPR4 antagonist treatment reduced CD4^+^ and CD8^+^ immune cell clusters in the lungs of SARS-CoV-2-infected K18-hACE2 mice. Black arrows indicate immune cell clusters. Scale bar = 20 μm.

**Supplementary Figure 5.**
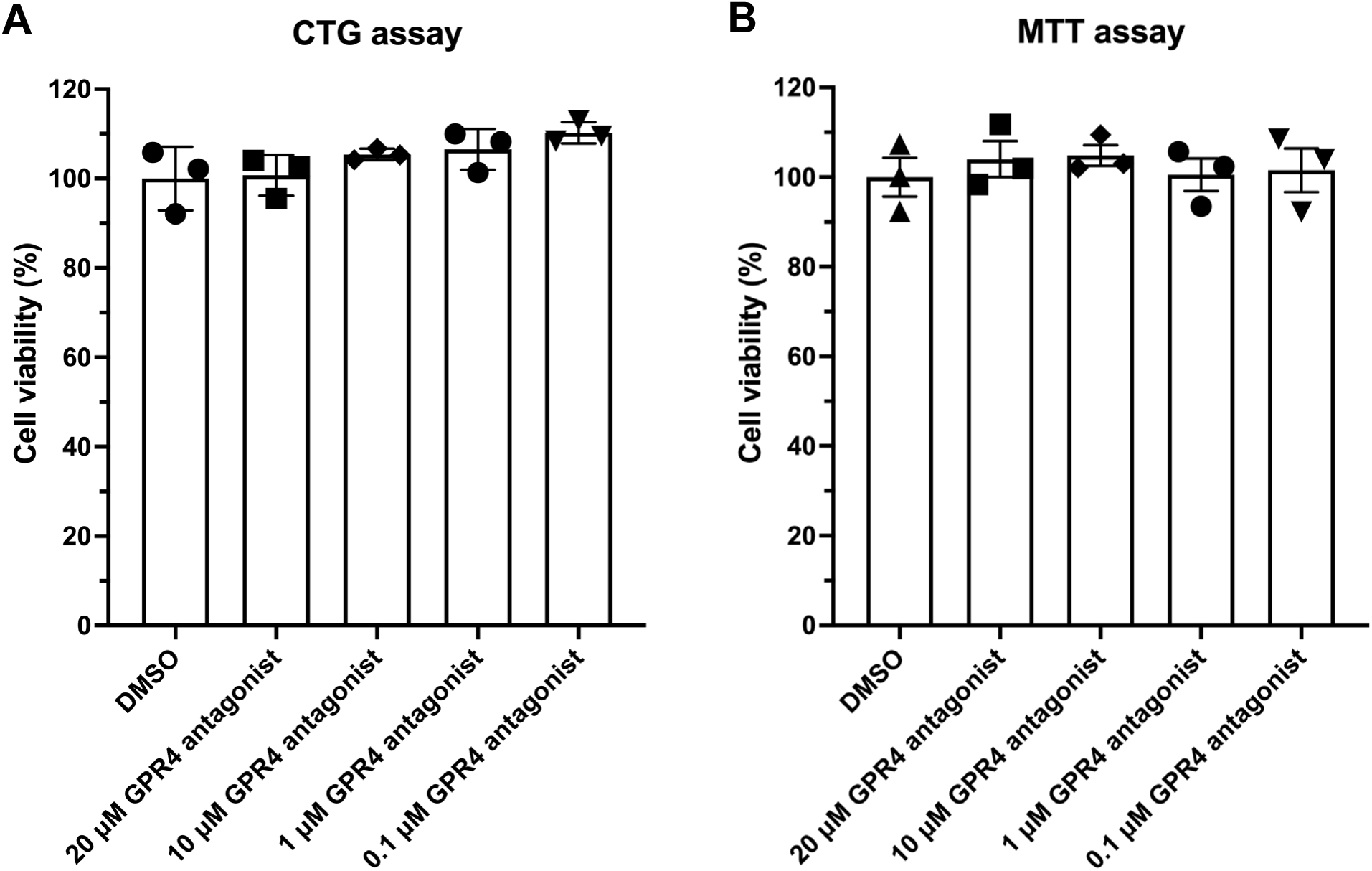
Viability of Vero E6 cells treated with the GPR4 antagonist. The viability of Vero E6 cells was approximately 100% relative to the DMSO control in the presence of 20 μM, 10 μM, 1 μM, and 0.1 μM GPR4 antagonist for 24h. (A) CellTiter-Glo (CTG) assay. (B) MTT assay.

**Supplementary Figure 6.**
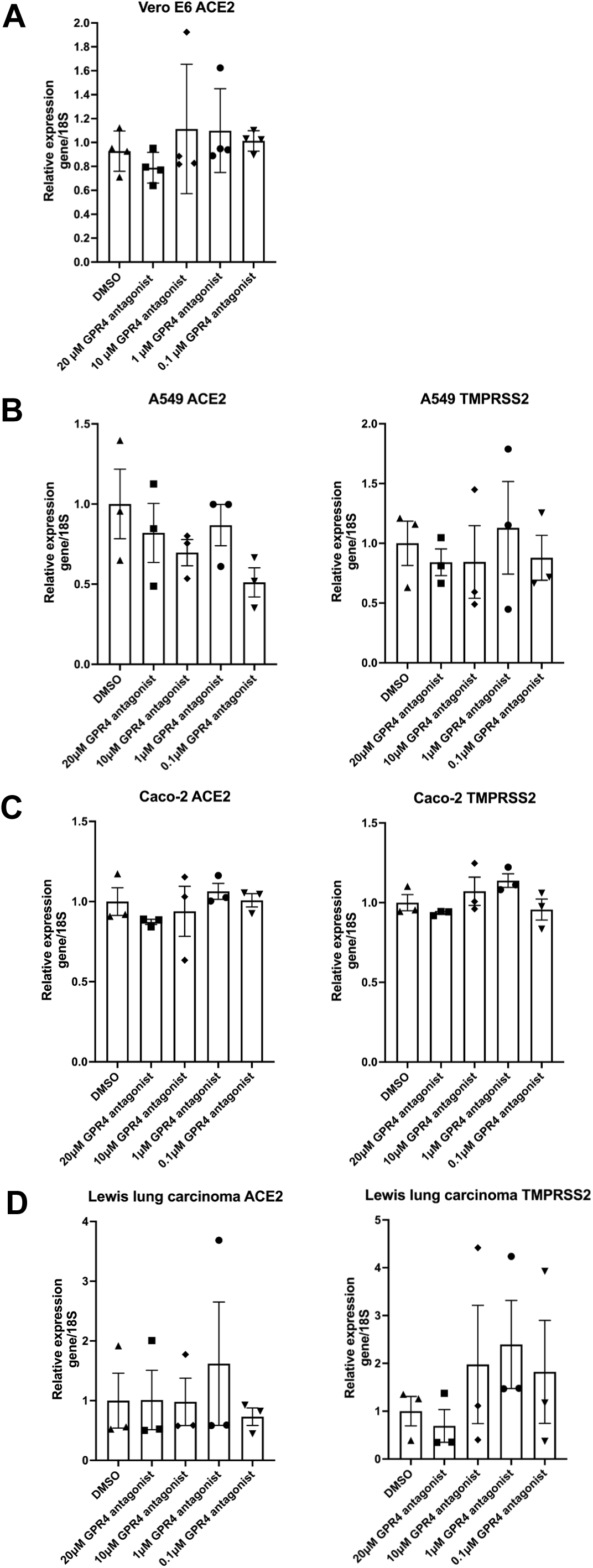
ACE2 and TMPRSS2 RNA expressions in cells treated with the GPR4 antagonists *in vitro*. Quantitative RT-PCR was performed to assess ACE2 or TMPRSS2 RNA expressions in Vero E6, A549, Caco-2, and Lewis lung carcinoma cells after a 24h GPR4 antagonist treatment. Two-tailed Student’s t-tests were used to compare ACE2 and TMPRSS2 expressions between vehicle and GPR4 antagonist treatment in cell lines.

